# The non-steroidal anti-inflammatory drug nimesulide kills *Gyps* vultures at concentrations found in the muscle of treated cattle

**DOI:** 10.1101/2021.08.31.457916

**Authors:** Toby. H. Galligan, Rhys. E. Green, Kerri. Wolter, Mark. A. Taggart, Neil. Duncan, John. W. Mallord, Dawn. Alderson, Yuan. Li, Vinny. Naidoo

## Abstract

Throughout South Asia, cattle are regularly treated with non-steroidal anti-inflammatory drugs (NSAIDs) and their carcasses are left for scavengers to consume. Residues of the NSAID diclofenac in cattle carcasses caused widespread mortality and catastrophic population declines in three species of *Gyps* vulture during the 1990s and 2000s. Diclofenac is now banned, but other NSAIDs are used in its place. Different lines of evidence, including safety testing in *Gyps* vultures, have shown that some of these other NSAIDs are toxic, or probably toxic, to vultures. The NSAID nimesulide is widely available and commonly used, and has been found in dead vultures with signs of renal failure (i.e. visceral gout) and without the presence of diclofenac and/or other vulture-toxic NSAIDs. Nimesulide is therefore probably toxic to vultures. Here, we report safety testing of nimesulide in *Gyps* vultures. In a controlled toxicity experiment, we gave two vultures the maximum likely exposure of nimesulide calculated from initial pharmacokinetic and residue experiments in cattle. Two other control birds were given an oral dose of water. Both vultures dosed with nimesulide died within 30 h, after showing outward signs of toxicity and increases in biochemical indicators of renal failure. Post-mortem examinations found extensive visceral gout in both vultures. Both control vultures survived without biochemical indicators of renal failure. With this evidence, we call for an immediate and comprehensive ban of nimesulide throughout South Asia to ensure the survival of the region’s Critically Endangered vultures. More generally, testing the impacts of drugs on non-target species should be the responsibility of the pharmaceutical industry, before their veterinary use is licensed.

## 1. Introduction

Pharmaceuticals in the environment can impact wildlife populations in a variety of lethal and sublethal ways (Arnold *et al.* 2014; Bean and Rattner 2018; Saaristo *et al.* 2018; Ulrika and Wong 2019). A catastrophic example of pharmaceutical pollution causing death and declines in wildlife populations is that of diclofenac and its impact on *Gyps* vultures in South Asia (Pain *et al.* 2008). *Gyps* vultures are highly sensitive to the non-steroidal anti-inflammatory drug (NSAID) diclofenac (Oaks *et al.* 2004; Shultz *et al.* 2004; Swan *et al.* 2006a). The widespread and frequent use of diclofenac to treat livestock, combined with the traditional practice of leaving cattle carcasses for vultures to consume in South Asia, caused catastrophic declines in three *Gyps* species (i.e., white-rumped vulture *G. bengalensis,* long-billed vulture *G. indicus* and slender-billed vulture *G. tenuirostris*; Prakash *et al.* 2007; Green *et al.* 2004, 2007). Hardest hit was the white-rumped vulture population in India, formerly one of the most abundant bird populations in the world, which declined by 99.9% between 1992 and 2007, a loss of an estimated 40 million birds (Prakash *et al.* 2007). Evidence suggests that other species of vultures and scavenging birds of prey are also intolerant to diclofenac and have suffered similar declines (Cuthbert *et al.* 2006; Acharya *et al.* 2009, 2010; Sharma *et al.* 2014). Five species of vultures, resident to South Asia, were uplisted to Critically Endangered or Endangered as a result of declines known or suspected to be caused by diclofenac (Birdlife 2017). Veterinary diclofenac is now banned throughout South Asia and declines in most vulture populations have slowed and in some have been reversed (Chaudhary *et al.* 2012; Galligan *et al.* 2014, 2019; Paudel *et al.* 2015; Prakash *et al.* 2017). However, the illegal use of human formulatons of diclofenac in veterinary care is still occurring, especially in India, albeit at lower levels than before the bans (Galligan *et al*. 2020). Furthermore, all vulture populations remain small and some are still declining (see Prakash *et al.* 2015; Paudel *et al.* 2016).

Diclofenac is not the only vulture-toxic NSAID used to treat cattle in South Asia. Recent surveys of pharmacies selling veterinary drugs in India, where the greatest variety of NSAIDs exists, found eleven active ingredients among products (Galligan *et al*. 2020). Furthermore, in a survey of cattle carcasses available to wild vultures in India, detectable NSAID residues were found in 16.2% of carcasses (RSPB/BNHS/ERI unpublished data). Only one NSAID, meloxicam, has been demonstrated to be safe to *Gyps* vultures (Swan *et al.* 2006b; Swarup *et al.* 2007); while four others – carprofen (Fourie *et al.* 2015; Naidoo *et al.* 2017), flunixin (Zorilla *et al.* 2014; Fourie *et al.* 2015; Herrero-Villar et al. 2020), ketoprofen (Naidoo *et al.* 2010b), and phenylbutazone (Fourie *et al.* 2015) – caused clinical signs of toxicity and/or death in *Gyps* vultures. Another NSAID, aceclofenac, rapidly metabolised into diclofenac in cattle (Galligan *et al.* 2016) and will, therefore have the same catastrophic effect on vultures. Nimesulide is the only NSAID to have ever been found in dead vultures in South Asia with visceral gout (a clinical sign of NSAID toxicity in vultures), and without diclofenac residues (Cuthbert *et al.* 2016, Nambirajan *et al*. In press), providing indirect evidence of the drug’s toxicity. Nimesulide is commonly sold in pharmacies for the treatment of cattle in both India (up to 37.3% of pharmacies sampled within the State of Haryana in 2017) and Nepal (up to 13.7% of pharmacies sampled within the western Terai in 2016) (Galligan *et al*. 2020). This highlights the importance and urgency for nimesulide to be safety tested in *Gyps* vultures, the results of which will inform conservation actions across South Asia.

Using the brand of nimesulide found throughout South Asia (‘Nimovet’), we report safety-testing of the drug in *Gyps* vultures. The safety test consisted of three experimental studies conducted in South Africa in domesticated cattle *Bos taurus* and wild-rescued *Gyps* vultures with serious injuries that prevented their release. First, we undertook a pharmacokinetic study of nimesulide in cattle to determine the time at which the concentration of the drug is at its greatest in plasma (*T_max_*). Second, we undertook a tissue residue study of nimesulide in cattle to determine the highest concentration of the drug among tissues at *T_max_* and used this to calculate the maximum likely exposure (*MLE*) of nimesulide in cattle tissue for *Gyps* vultures at a single feeding. Third, we undertook a toxicity study of the *MLE* of nimesulide in cattle tissue in individual *Gyps* vultures to determine the toxicity of the drug at the worst-case scenario. We chose not to conduct safety testing at lower doses of the drug to minimise the number of birds exposed to potential harm, both physical and psychological, from the experimental process.

## 2. Methods

### 2.1 Research permission

We were permitted to conduct this study by the Research Committee and Ethics Advisory Committee (EAC2016-02) of the RSPB Centre for Conservation Science, RSPB, UK, and the Research Committee of the Faculty of Veterinarian Sciences, University of Pretoria, and the Animal Ethics Committee of the University of Pretoria (V031/13), South Africa. We were permitted to work on an Endangered species (*Gyps coprotheres*) by the South African Department of Environment Affairs (Permit S02655). CITES South Africa (permit no: 206268) and the Medicines Control Council of South Africa (27/2/2 VCT/12/2016) granted us permits to export vulture and cattle samples from South Africa; while CITES UK (584542/01) and the Animal Plant Health Agency, Department of Environment, Food and Rural Affairs, UK, (ITIMP19.0188) granted us permits to import vulture and cattle tissue samples to the UK.

### 2.2 Cattle test subjects

We used *Bos taurus* cattle in each experiment. Specifically, four (identifier codes: C1-C4) Friesian cows at 9 months of age in the pharmacokinetics study and four (C5-C8) Nguni cows at 9 months of age in the tissue residue study. We acquired cattle from commercial cattle farmers in Pretoria, South Africa. Cattle were housed in non-quarantine outdoor camps at the University of Pretoria for one month to ensure they were free of veterinary drugs, according to established clearance times. One week before experimentation, we moved the cattle to temperature-controlled stables at the Biomedical Research Centre, University of Pretoria. We provided food, water and exercise daily to the cattle in both locations; and bedding in the stables. One day before experimentation, we gave each cow a complete veterinary examination and deemed them fit and healthy. We monitored the cattle throughout the experiment for signs of adverse reactions to nimesulide but observed none. The cattle we used in the pharmacokinetics study were returned to their owners with an enforced three-month withdrawal period. The cattle we used in the tissue residue study were euthanized at the facility.

### 2.3 Vulture test subjects

We safety tested nimesulide in Cape vultures (*Gyps coprotheres*): we used two immature (identifier codes: G32745 and G34994) and two mature (G32746 and G32753) vultures. This species is known to be sensitive to two other NSAIDs: ketoprofen (Naidoo *et al.* 2010b); and carprofen (Naidoo *et al.* 2017). Each vulture was found injured in the wild. We rescued and treated them at VulPro, South Africa, following standard protocols and with the intent of releasing them when they were deemed fit and healthy enough to survive in the wild. However, these vultures had serious injuries that made them non-releasable, specifically: G32745 had one broken leg and one dislocated leg, which made standing and walking impossible; G32746 had a dislocated leg, which made standing and walking impossible; G32753 had a missing hind toe and severe bumblefoot, which made standing and walking difficult; and G34994 had had a severely broken wing amputated, which made flying impossible. G32745 had been rescued days before the toxicity experiment and therefore was only kept in our small hospital aviaries (dimensions [h x l x w]: 3 x 3 x 6 m). The other three vultures had been rescued several months before the toxicity experiment and had been moved to larger communal aviaries (at least 6 x 8 x 48 m). A day before experimentation, we moved the vultures in the communal aviaries to the hospital aviaries. In the communal aviaries, vultures were given food (whole pig carcasses), water, shade, perches and room for short flapping flight. In the hospital aviaries, vultures were given food, water, shade, perches and a small shed for shelter. We fed all vultures following a standard regime but stopped feeding two days before the first day of the toxicity experiment. One day before the first day of the toxicity experiment, we gave each vulture a complete veterinary examination and deemed them fit and healthy aside from their injuries described above. We resumed feeding with a small meal (i.e., 250 g of mixed pig tissues) one day after the first day of the toxicity experiment. At the end of the toxicity study, we returned surviving vultures to large communal aviaries and standard feeding regimes (Wolter, Neser and Hirschauer 2015).

### 2.4 Pharmacokinetics experiment in cattle

We weighed each cow (n = 4, mean ± SD = 209.25 ± 7.63 kg). Next, we collected a 10-ml sample of blood from the jugular vein of each cow into an evacuated heparinised tube. Then, we treated each cow once with an injectable formulation of nimesulide (Nimovet, Indian Immunologicals Ltd., India) intramuscularly at the standard dose of 2 mg kg^−1^ bw. Following this dosing, we collected blood samples from each cow into evacuated heparin tubes at 0.25, 0.75, 1.50, 2.00, 3.00, 5.00, 7.00, 9.00, 12.00, 24.00, 36.00, 48.00, 96.00 and 120.00 h after dosing. We centrifuged all samples within 2 h at ~3000 *g* for 15 min, and transferred the supernatant (plasma) into plastic screw-top vials and stored the vials at −80 °C.

### 2.5 Tissue residue experiment in cattle

We undertook this second experiment once we had obtained data from the first experiment (see method below). Approximately two months separated these experiments. We weighed each cow (n = 4, mean ± SD = 175.50 ± 20.14 kg). We then treated them intramuscularly with double the recommended dose of nimesulide (Nimovet; i.e., 4 mg kg^−1^ body weight (bw)) once a day for three days. We doubled the dose to simulate the apparent behaviour of veterinarians and livestock owners in South Asia, based on measured NSAID residues in the tissues of dead cattle that show that these animals sometimes received much more than the recommended dose of various drugs (Taggart *et al.* 2007), especially as an animal’s ailment progresses towards the end of its life. We gave all injections on one side of the neck of the cattle. We slaughtered the cattle at *T_max_* using a captive bolt and pithing. We harvested muscle from the injection site at the side of the neck and muscle from the hindquarters (hereafter, non-injection site muscle). We also harvested the liver and one kidney from each cow. We took three replicate samples from the injection site muscle and a single sample from each of the other tissues. We stored samples and harvested tissues at −80 °C.

### 2.6 Calculating the MLE and dose of nimesulide for individual vultures

First, we calculated the mean daily energy use (*DEU* in kJ d^−1^) for individual vultures as *DEU* = 668.4\**W_i_*^0.622^ (Mundy *et al*. 1992), where *W_i_* is the bodyweight for the individual vulture *i.* Second, we calculated the maximum likely meal (*M_i_* in kg) for individual vultures as *M_i_ = 3*\*(*DEU*/*EG*), where *EG* is the content of assimilable energy of ungulate tissue (5160 kJ kg^−1^), which is the product of its energy content (6000 kJ kg^−1^) and the proportion of ingested energy that is assimilated by a vulture (0.86). Observations of Cape vultures suggests that the maximum meal size of *Gyps* vultures is typically about three times the maximum amount of food required per day (S. Piper, quoted in Swan *et al*. (2006a), Supplementary Information, Protocol S1). If the mean weight of the tissue with the highest mean concentration of nimesulide (*V_max_*) was greater than *M_i_*, we calculated the *MLE* for individual vultures as *MLE* = *R_max_ *M_i_*, where *R_max_* is the mean concentration of nimesulide in the tissue with the highest mean concentration. If *V_max_* was less than *M_i_*, we calculated the maximum likely exposure (*MLE* in mg) for individual vultures as *MLE* = *R_max_*V_max_* + *R_next_*(M_i_-V_max_*), where *R_next_* is the mean concentration of nimesulide in the tissue with the second highest mean concentration. We calculated the dose in mg kg^−1^ (*D_1_*) as *D_1_* = *MLE/W_i_* for individual vultures and ml (*D_2_*) as *D_2_ = (MLE*W_i_*)/*U* for individual vultures, where *U* was the concentration of nimesulide (Nimovet) used (100 mg ml^−1^).

### 2.7 Toxicity experiment in *Gyps* vultures

We undertook this third experiment once we had obtained data from the second experiment (see method above and below). Approximately four months separated these experiments. We assigned G32745 and G32746 to the treatment group and G32753 and G34994 to the control group. We did this non-randomly because the injuries of G32745 and G32746 impacted more on their lives than those of the other two vultures and we had planned to euthanize them irrespective of the outcome of the toxicity experiment. As a result, each treatment group consisted of one mature and one immature bird. Before the first blood sampling, birds were deemed to be clinically healthy, with the exception of their particular injury. We began the toxicity experiment by weighing each vulture and collected a baseline blood sample. We fitted a catheter into the tarsal vein of three vultures (G32745, G32746 and G34994) to collect blood, but were unable to fit a catheter to the remaining vulture; hence, we used a syringe to collect blood from that vulture (G32753). To facilitate the flow of blood in vultures with catheters, each time we sampled blood, we first injected 1 ml of heparinised saline and immediately drew and discarded 1 ml of blood; we then collected 5 ml of blood and injected another 1 ml of heparinised saline. Blood was collected into heparinised tubes. We then gave the treatment group a dose of injectable nimesulide (Nimovet) based on the *MLE* of nimesulide for a vulture of their body weight (17.58 mg/kg body weight (bw)). The drug was given orally via a syringe followed by 2 ml of water. We gave the control group an oral dose of 5 ml of water. We then sampled 5 ml of blood into heparinised tubes as above from each vulture at 2, 6, 12, 24 and 48 h after dosing. We centrifuged all blood samples within 15 minutes at ~3300 *g* for 15 min and transferred the plasma (supernatant) into two plastic screw-top vials. Vials were initially stored in a standard freezer but were transferred to a −80 °C freezer within 30 h. We observed vultures regularly throughout daylight hours (05:00-18.30) for 7 days after dosing and recorded any abnormal behaviour. Our specialist veterinary pathologist (N. Duncan, Department of Paraclinical Sciences, Faculty of Veterinary Science, University of Pretoria) performed post mortem examinations on all vultures that died.

### 2.8 Analysing samples for nimesulide concentration

We transported the plasma and tissue samples from cattle and one set of plasma samples from vultures on dry ice to SAC Veterinary Consulting Services, Scotland’s Rural College (SRUC), UK. We extracted 0.3-ml subsamples from each plasma sample and three 0.5-g subsamples from each tissue sample into 3 ml of HPLC grade acetonitrile. For plasma samples, we vortex mixed for 20 s, rested at room temperature for 600 s and vortex mixed again for another 20 s, before centrifuging. For tissue, we homogenised with a VDI 12 (VWR) homogeniser, before centrifuging. In both cases, we centrifuged at 2000 rpm, 671 *g* for 600 s, and transferred the supernatant directly into amber 2-ml screw-top LC vials using a syringe filter (0.2 μm HDPE disposable) and stored the supernatant at −20°C. We analysed extracts for nimesulide concentration at the Environmental Research Institute (ERI), University of the Highlands and Islands, UK. Nimesulide concentrations were determined using liquid chromatography-electrospray ionisation triple quadrupole mass spectrometry (LC-ESI-MS/MS) utilising a methodology adapted after Taggart et al. (2009). Nimesulide was detected in negative ion mode utilising a parent target mass of 307 m z^−1^ and two daughter ions of 229 m z^−1^ (quantitation) and 198 m z^−1^ (confirmation) in multiple reaction monitoring (MRM) mode. Mean recovery of nimesulide spiked into blank plasma (n = 8) and liver tissue (n = 6) at two different concentration levels and extracted as above was 89.2% and 140.0% respectively. Final concentrations were calculated following correction for these extract recovery levels. The limit of quantification for the analysis was 4 ng ml^−1^ and 10 ng g^−1^ for plasma and tissue respectively.

### 2.9 Pharmacokinetic evaluation

Pharmacokinetic parameters for vulture plasma samples were ascertained by non-compartmental modelling in Kinetica 5.1 (Thermo 2012). The maximum plasma concentration (*C_max_*) and the time to reach it (*T_max_*) were read from the plasma concentration versus time profile. The last quantifiable time point *C_last_* and the linear trapezoidal rule was used to calculate the area under the curve (*AUC_last_*) and the area under the moment curve (*AUMC_last_*) as (*AUC_last_* = Ʃ([*T_last_*-*T_last_*-1]*[*C_last_*+*C_last_*-1]/2)) and (*AUMC_last_* = Ʃ([*T_last_*-*T_last_*-1]*[*T_last_*\**C_last_*+*T_last_*-1\**C_last_*-1]/2)), respectively. The elimination rate constant (*λ*) was calculated by ordinary least squares regression of the terminal three points of the curve after natural logarithmic transformation; and subsequently, the half-life of elimination (*T_half_*) was calculated as ln(2)/*λ*. The mean residence time (*MRT*) was calculated as *AUMC_last_*/*AUC_last_* and the area under the curve to infinity (*AUC_inf_*) was calculated as *AUC_last_* + *C_last_*/*λ*. For nimesulide administered intramuscularly and orally, the apparent volume of distribution (*V_z_*/*F*) was calculated as dose/(*AUC_last_*\**λ*), the apparent volume of distribution at steady state (*V_ss_*/*F*) was calculated as (dose\**MRT*)/*AUC_last_* and the apparent clearance (*Cl*/*F*) was calculated as dose/*AUC_last_*; and for nimesulide administered intravenously, the actual volume of distribution (*V_z_*), actual volume of distribution at steady state (*V_ss_*) and actual clearance (*Cl*) were calculated by first finding the fraction of absorption (*F*) and dividing this by the apparent measures of these parameters. *F* was calculated as a bird’s extravascular *AUC_inf_* divided by the pooled *AUC_inf_* from the intramuscular profile. All parameters are presented as geometric means with standard deviations.

### 2.10 Analysing samples and vultures for clinical pathology

The second set of plasma samples from vultures were analysed by the Veterinary Clinical Pathology Laboratory, University of Pretoria. Standard analytes were measured using a Cobas Integra 400. We were particularly interested in the changes in concentrations of alanine transferase (ALT), alkaline phosphatase (ALP), potassium, sodium, calcium, urea and uric acid, all of which are known indicators of renal failure (Naidoo *et al.* 2017). Change in analyte concentrations was considered important if a value at a given time was different to the 1) baseline value, 2) previous measured value, and 3) higher or lower than the normal range of values delineated by the highest and lowest value among those for the control group at any time during the series.

### 2.11 Post-mortem examination of vultures

We performed a post-mortem examination on all vultures that died during the toxicity test. We began with a terminal blood sample collected, processed and analysed as described above. We performed a necropsy, focussing on the viscera since visceral gout – the accumulation of urates on the viscera – was associated with toxic death in *Gyps* vultures during safety testing of diclofenac (Oaks *et al.* 2004; Swan *et al.* 2006a), ketoprofen (Naidoo *et al.* 2010b) and carprofen (Naidoo *et al.* 2017). Visceral gout in birds results from renal failure. *Gyps* vultures show zero-order elimination after exposure to some NSAIDs, which suggests that they have limited enzyme capacity for processing these drugs (Naidoo *et al.* 2010a; Naidoo *et al.* 2017). An inability to process some NSAIDs leads to renal failure and presents as visceral gout on necropsy. Tissue samples were collected during the necropsy, fixed in 10% buffered formalin and routinely processed, embedded in paraffin wax and cut at 4 μm. After mounting, the sections were stained routinely with Haematoxylin and Eosin (H&E).

## 3. Results

### 3.1 Pharmacokinetics experiment in cattle

Plasma concentrations versus time profiles among cattle C1-C4 were consistent. The greatest concentrations of nimesulide in individual cattle ranged between 25.68 and 34.80 μg ml^−1^. The greatest mean concentration among cattle (*C_max_*) was 28.90 μg ml^−1^, which was measured at 3 h after dosing; therefore, *T_max_* was 3 h (Supplementary Figure 1).

### 3.2 Tissue residue experiment in cattle

The tissue with the highest mean concentration of nimesulide *R_max_* among cattle C5-C8 was the injection site muscle at 134.08 mg kg^−1^ (Supplementary Table 1), and its mean weight, *V_max_,* was 1.11 kg (Supplementary Table 2).

### 3.3 Calculating the MLE and dose of nimesulide for individual vultures

The two vultures to be treated with nimesulide shared a *W_i_* of 8.5 kg and therefore their *M_i_* was 1.47 kg. Since this *M_i_* was greater than *V_max_*, we calculated the *MLE* as *R_max_*V_max_* + *R_next_*(M_i_-V_max_*), where *R_next_* was the concentration of the non-injection site muscle at 0.945 mg kg^−1^ (Supplementary Table 1). We calculated an *MLE* of 149.44 mg kg^−1^, a *D_1_* of 17.58 mg kg^−1^ and a *D_2_* of 1.49 ml (rounded to 1.50 ml) for both G32745 and G32746.

### 3.4 *Toxicity experiment in* Gyps *vultures*

Vulture G32745 showed no immediate adverse reaction to nimesulide, whereas vulture G32746 attempted to spit out the drug. G32746 may have received a lower unknown dose as it was observed expelling liquid, although C_*max*_ was similar for both birds (see below), so G32746 likely received a substantial amount of the dose. From 3-4 h after dosing until 25-26 h after dosing, both G32745 and G32746 showed relief from pain and were observed standing and walking. Beyond 25-26 h after dosing, both vultures showed signs of toxicity. G32746 lay awkwardly on its side with its wings spread, head and neck often on the ground and eyes often shut. G32745 lay awkwardly on its front with wings spread, but head and neck raised and eyes open. Both vultures showed laboured breathing, raised rectal lumps and spayed regimes. G32745 died 27.50 h after dosing, despite appearing less effected by nimesulide than G32746, which died 29.34 h after dosing. The control group (G32753 and 34994), remained alert and active until the experiment was terminated, 48 h after dosing.

### 3.6 Analysing samples for nimesulide concentration

Pharmacokinetic profiles (Figure 1) and parameters (Table 1) were consistent between the two vultures treated with nimesulide. Both vultures showed rapid absorption in the first six hours, followed by a slower elimination over the next 18-23 hours before their deaths (Figure 1). They shared a T_max_ value and had similar C_max_ values (Table 1). Vulture 32746 showed a greater extent of absorption and faster elimination than 32745 (Figure 1), which was reflected in several parameters associated with the area under the curve, mean residence time and elimination (Table 1). The terminal concentration in 32756 (and probably 32745) was the lowest after dosing (Figure 1).

**Figure 1:**
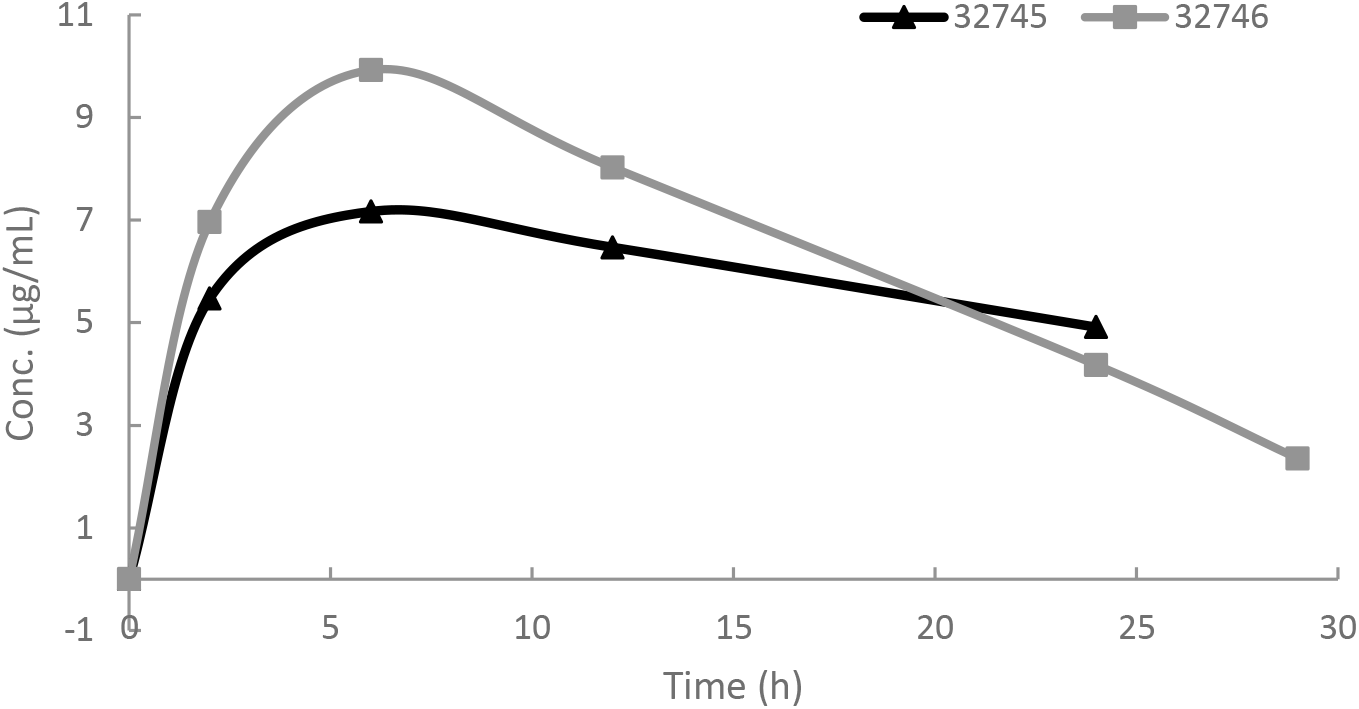
Pharmacokinetic profiles for the two vultures *Gyps coprotheres* (32745 and 32746) treated with nimesulide at 17.58 mg/kg bw by the oral route.

**Table 1:** Pharmacokinetic parameters for the two vultures *Gyps coprotheres* (32745 and 32746) treated with nimesulide at 17.58 mg/kg bw by the oral route.

| Parameter | Units | Vulture |  | Mean | SD | CV (%) |
| --- | --- | --- | --- | --- | --- | --- |
|  |  | 32745 | 32746 |  |  |  |
| $C_{max}$ | µg/mL | 7.00 | 9.00 | 7.94 | 1.41 | 0.18 |
| $T_{max}$ | h | 6.00 | 6.00 | 6.00 | 0.00 | 0.00 |
| $AUC_{last}$ | µg/mL*h | 128.00 | 174.00 | 149.24 | 32.53 | 0.22 |
| $AUC_{extra}$ | µg/mL*h | 127.97 | 29.42 | 61.36 | 69.68 | 1.14 |
| $AUC_{tot}$ | µg/mL*h | 255.97 | 203.43 | 228.19 | 37.15 | 0.16 |
| $\%AUC_{extra}$ | % | 49.99 | 14.46 | 26.89 | 25.12 | 0.93 |
| $\lambda$ | 1/h | 0.03 | 0.08 | 0.05 | 0.03 | 0.66 |
| $AUMC_{last}$ | µg/mL*(h)<sup>2</sup> | 1464.00 | 2131.00 | 1766.29 | 471.64 | 0.27 |
| $T_{half}$ | h | 22.02 | 8.98 | 14.06 | 9.22 | 0.66 |
| $MRT$ | h | 33.60 | 16.54 | 23.58 | 12.06 | 0.51 |
| $Cl$ | L/h*kg | 0.07 | 0.08 | 0.07 | 0.01 | 0.16 |
| $V_z$ | L/kg | 2.11 | 1.08 | 1.51 | 0.73 | 0.48 |
| $V_{ss}$ | L/kg | 2.23 | 1.38 | 1.76 | 0.60 | 0.34 |

### 3.6 Analysing samples for clinical pathology

The two treated vultures showed increases in plasma uric acid and potassium concentrations above the normal ranges for each analyte (Figure 2). Both vultures showed uric acid concentrations above our maximum level of detection (15 mmol/L) at 24 h, which represented a minimum 27-79-fold increase over their baseline values. At 24 h, potassium concentrations showed a 2-4-fold increase over baseline values for each vulture. The treated vultures also showed slight decreases in plasma calcium concentrations (Figure 2). One treated vulture (G32745) showed an increase in ALT (which, at 24 h, was 5-times greater than the baseline value), consistently high concentrations of ALP and consistently low concentrations of sodium; whereas, the other treated vulture (G32746) showed no change outside the normal range for ALT, ALP and sodium. At the time of death, G32746 plasma showed its highest potassium concentration, its lowest concentration of calcium and a level of uric acid slightly lower than at 24 h (Figure 2).

**Figure 2:**
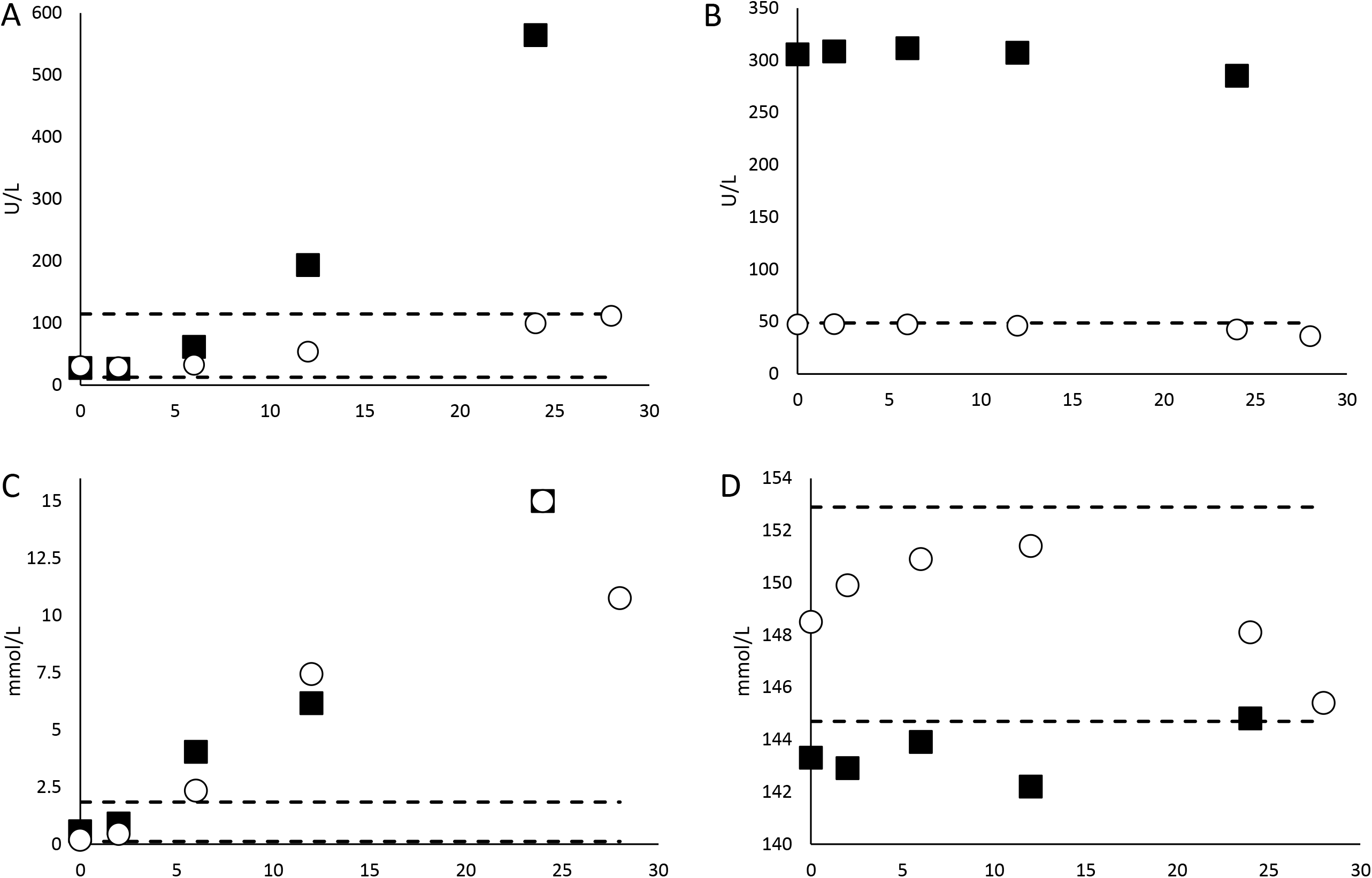
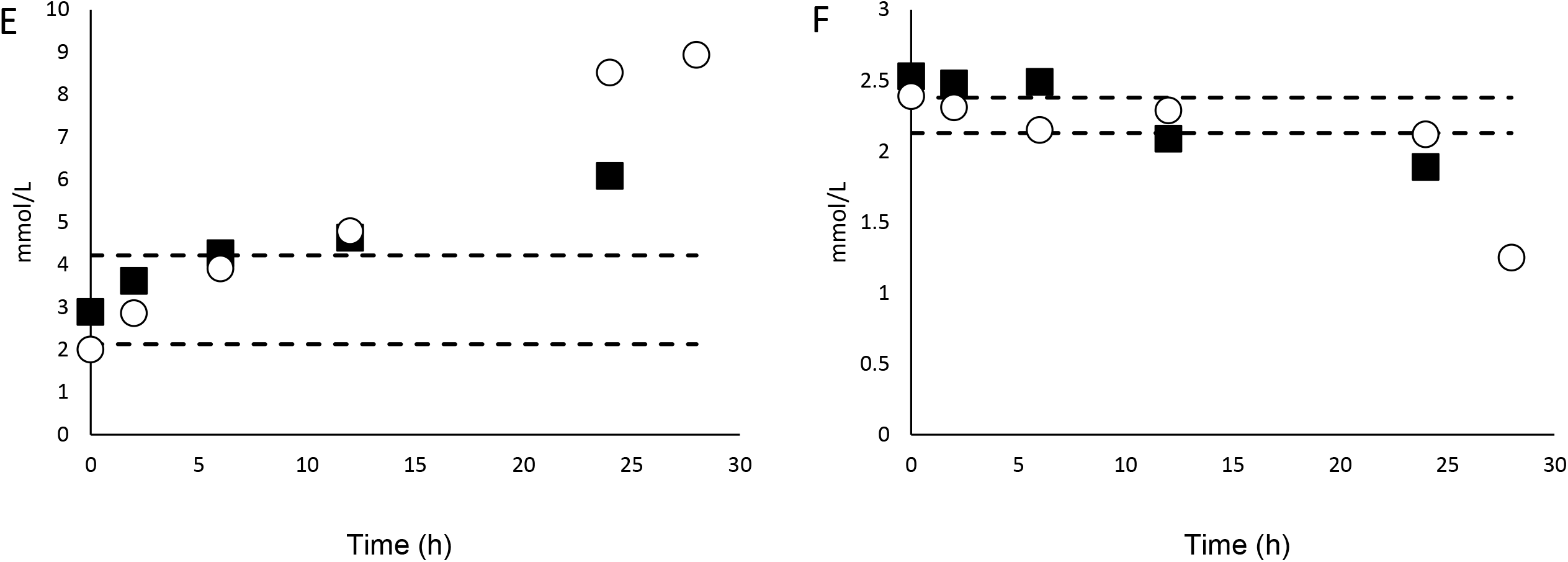
Change over time in plasma ALT (A), ALP (B), uric acid (C), sodium (D), potassium (E) and calcium (F) concentrations in two vultures (solid squares = G32745 and open circles = G32746) treated with nimesulide at 17.58 mg/kg bw. G32746 likely received a lower unknown dose as it was observed expelling liquid immediately after dosing. Dashed lines show the normal range of concentrations for each analyte, delineated by the lowest and highest measurements taken from two control vultures.

### 3.7 Post-mortem examination of vultures

A terminal blood sample was obtained from G32746 at 28.09 h after dosing, but not from G32745. Post mortem examination found visceral gout in both vultures in the form of small and scattered tophi (i.e., deposits of crystalline uric acid) in the kidneys, spleen, lungs and liver. Histopathology found several lesions in association with these tophi in both vultures; and necrosis in the form of pyknosis, karyorrhexis, and desquamation was evident in both vultures (Figure 3). While G32746 also showed evidence of severe necrotising tracheitis and bacterial emboli in the brain at necropsy, subsequent histopathological evaluation confirmed signs of only ulcerative tracheitis with adherent fibrin and bacteria with no evidence of CNS damage. The damaged trachea would have been the entry point for the bacteria, but as there were no signs of bacterial emboli within the kidney, nor any changes cosnsistent with inflammation within the kidnesy, it is unlikely that it affected any of the other parameters. The cause of the tracheitis was considered incidental and not invested further, since we were confident of the cause of death.

**Figure 3:**
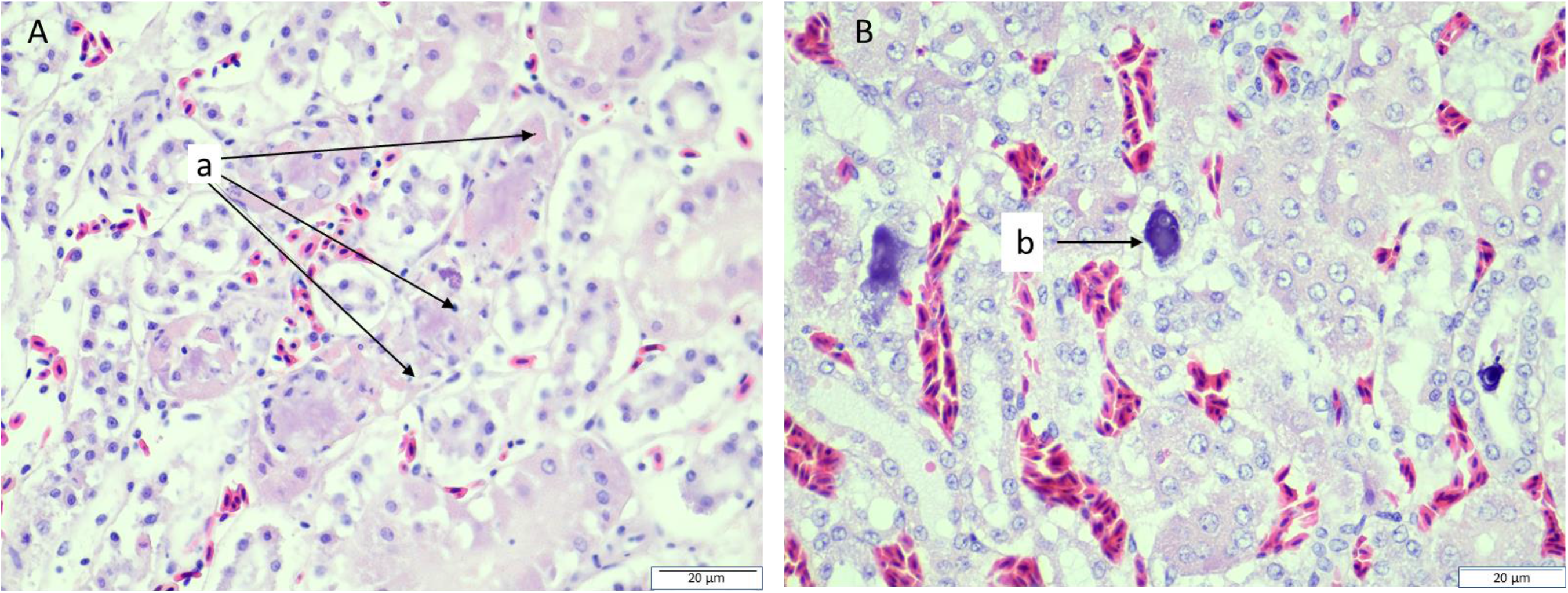
Histopathological images of kidney tissue from two Cape vultures (*Gyps coprotheres*; A = G32745 and B = G32746) treated with nimesulide at 17.58 mg/kg bw. Image A: increased cytoplasmic eosinophilia, nuclear pyknosis and karyorrhexis and desquamation from the basement membrane of the renal tubule cells. There are several renal tubules (a) containing dead and dying cells, with bright pink cytoplasm, the nucleus is either small and black (pyknosis) or consists of varying sized black dots (karyorrhexis). They are also no longer attached to the side of the tubule (desquamation), which occurs after death. Image B: centrally, a small tophus consisting of an area of necrosis with loss of normal architecture, being replaced by cell debris and spicules of uric acid. The purple blob (b) is a form of urate, called globular urate, which also occurs in cases of visceral gout.

## 4. Discussion

Our study supports the contention of Cuthbert *et al.* (2016) that nimesulide in cattle carcasses is killing *Gyps* vultures in South Asia. We safety tested nimesulide in an African *Gyps* species that is an appropriate surrogate test species for South Asia’s Critically Endangered *Gyps* species (Swan et al. 2006a). We gave nimesulide directly by the oral route, at twice the recommended dose, a level that is likely encountered by *Gyps* vultures in the wild, given that ailing cattle can receive progressively larger doses when close to death. The vultures absorbed the drug rapidly but eliminated it slowly. Nimesulide provided pain relief to the vultures throughout most of the absorption and elimination periods; however, uric acid started to accumulate with the onset of the elimination period and reached extremely high levels well before the end of the elimination period. This accumulation of uric acid killed the vultures as tophi affected numerous organs (i.e., visceral gout) causing widespread lesions and high concentrations of potassium in plasma is consistent with blood chemistry changes associated with cardiac failure (Sturkie 1986, Zandvliet 2005) and low concentrations of calcium in plasma likely indicated renal failure (Lierz 2003). Toxicity was only apparent in the vultures’ behaviour hours before death when concentrations of uric acid in plasma were extremely high. Behavioural, haematological, and histopathological evidence for nimesulide toxicity in *Gyps* vultures was similar to that for diclofenac (Oaks et al. Meteyer et al. 2005) and carprofen toxicity (Naidoo et al. 2017) in *Gyps* vultures. The vultures died with hours of each other, but G32745, an immature vulture, showed a slower rate of elimination than G32746, a mature vulture. Interestingly, G32745 also showed further signs of liver damage in tissue as well as in plasma. The increase in plasma ALP at the predisposed point (5 min before treatment) would indicate a pre-existing disease was present. Given its substantially high values, this is most likely linked to its known bone injury, as this is a known cause of substantial increases in serum ALP activity (Harr 2002). We did not test either vulture for infections or disorders that may have caused such pre-existing disease. However, the damage within the liver was just that of scattered tophi and necrosis. The hepatocytes themselves, as well as the bile ducts, did not show any changes consistent with exposure to hepatotoxins or viral / bacterial agents. Also, vulture G32746 spat out an unknown amount of nimesulide and the differences in response between the two vultures may be a result, at least in part, due to a difference in actual doses between vultures. Despite these differences that may have had an effect on finer details of their deaths, both vultures died in little more than a day after exposure to nimesulide at a level that is likely encountered by *Gyps* vultures in the wild.

In cattle, nimesulide was detected in the circulatory system not long after treatment; however, much of the drug remained at the injection site, resulting in a larger concentration in this muscle than in organs of the urinary system. There was also individual variation in drug absorption among cattle. Similar results were obtained for the NSAID, carprofen, in a similarly designed experiment to safety-test that drug in *Gyps* vultures (Naidoo et al. 2017). In both studies, we calculated the worst-case scenario dose for vultures based on the concentrations present at the injection site. Whereas only one of two vultures died when given a dose of carprofen, both vultures died when given nimesulide. However, the sample of vultures used in this study is too small to conclude that individual variation doesn’t exist in the population. Safety testing of carprofen also showed that doses of the drug based on (lower) concentrations found in organs did not kill vultures (although, once again, a small sample conceals possible individual variation). The present study was designed to evaluate the worst-case scenario, therefore we are confident that a bird feeding on tissues from the injection site will succumb to toxicity. Similarly high NSAID concentrations have been found in dead vultures (e.g., flunixin, Zorrilla *et al*. 2014), suggesting some birds are exposed to elevated drug levels through consuming the carcass of a recently-treated animal and/or tissues with high drug concentrations. However, our study cannot identify the level of toxicity that may result from the consumption of lower concentrations of the drug found in other areas of the carcass, as we are yet to establish the lower threshold for toxicity, i.e., the median lethal dose (LD50) may be lower than the concentrations evaluated in this study. From general toxicity principles, the LD50 tends to be determined by an individual’s inherent hepatic metabolising capacity and the rate of drug elimination. For another NSAID, ketoprofen, there was marked variation between individuals, with toxicity only being seen in those birds where zero order metabolism has been reached (Naidoo *et al*. 2010a). To clarify this further would thus require a larger sample size; however, we considered subjecting a larger sample of an endangered vulture speceis to lethal, as well as sub-lethal, but harmful, concentrations of nimesulide and the physical and psychological stress of the experimental process unethical and unnecessary. Our objective was to test whether nimesulide could kill vultures at exposure levels that could be encountered in the wild, and we have shown that it can. Dead wild *Gyps* vultures have previously been found in India with nimesulide residues and signs of renal failure (Cuthbert *et al* 2016, Nambirajan *et al*. 2021); we provide experimental confirmation of this NSAID’s toxicity to vultures.

Nimesulide is metabolised quickly in cattle,therefore treated individuals would have to die shortly after administration to maintain the high concentrations observed in this study. However, this is not impossible, because NSAIDs are often given daily as part of palliative care for dying cattle in South Asia (V Prakash pers. comm.). In addition, metabolism in dying cattle is most likely to be slower than in young and healthy cattle that were treated in this study; thereby, dying cattle likely show slower elimination and thereby greater periods at high concentrations. We used double the recommended dose of nimesulide here, but again this reflects the use of other NSAIDs in South Asia (Taggart *et al.* 2009), which might also be linked to palliative care of dying cattle. While we did not feed vultures nimesulide-rich cattle tissue, but administered the drug orally, thus controlling for the route of absorption in vultures. We acknowledge that it is possible that vultures absorb nimesulide in a liquid at a faster rate than nimesulide in a solid (muscle) as a result of crop retention. However, this is unlikely to interfere much with overall exposure to the drug as the crop would only delay food entering the stomach by a few hours. While the impact of food on NSAID absorption in vultures is yet to be established due to the complexities of regurgitation as a natural defence after feeding, evidence from human studies has shown no major impact of meals on NSAID pharmacokinetics with absorption being delayed by a few hours (Moore *et al*. 2015). Thus while uncertainty exists in aspects of our experimental design, we are satisfied that we have simulated a realistic scenario for nimesulide exposure to vultures in South Asia.

Nimesulide concentrations in cattle liver were extremely low at the same time after dosing that it was extremely high in plasma; while it was high to extremely high in muscle, particularly close to the injection site. Drug concentrations in the renal organs are not expected to increase during the drug’s elimination phase. As a result, detecting nimesulide in the liver of dead cattle in the field is expected to be difficult, which supports the idea that the cattle carcass surveys for NSAIDs to date that sampled liver tissue only do not detect nimesulide despite its widespread availability and frequent sales from pharmacies selling drugs for animals (Galligan et al. 2020). Whether this phenomenon is unique to nimesulide is unknown. Carprofen is the only other NSAID that has been subject to a tissue residue experiment in cattle, which found the drug in similar concentrations in the liver, kidneys and non-injection site muscle at *T_max_* (Naidoo et al. 2017). Despite this, carprofen concentration, like nimesulide, is highest in muscle close to the injection site (Naidoo et al. 2017). For these reasons, future cattle carcass surveys should sample muscle from either side of the neck in addition to liver tissue to obtain more accurate data on a wider range of NSAIDs used to treat cattle in South Asia. As a consequence, great caution should be taken in interpreting the prevalence of nimesulide from current published and unpublished data from cattle carcass surveys that did not test for nimesulide in injection site muscle.

Like all NSAIDs, nimesulide inhibits the enzyme cyclo-oxygenase (COX) and thereby the synthesis of prostaglandins (Bennett and Vila 2000). More specifically, nimesulide is a COX-2 inhibitor, which better targets pain and inflammation and thereby causes fewer gastrointestinal complications (Suleyman et al. 2008). Nimesulide has analgesic, antipyretic and anti-inflammatory properties (Bennett and Vila 2000). The vulture-safe NSAID meloxicam is also a COX-2 inhibitor and confers similar benefits to treated animals (e.g. European Medicines Agency 2018). Meloxicam also outperforms nimesulide in its effectiveness and relative safety to the target animals (usually mammalian livestock and pets; SAVE 2016). A review of experimental and clinical studies reporting effects of nimesulide and meloxicam treatment in non-human mammals, found 117 studies for meloxicam and only one study for nimesulide (SAVE 2016). The single study examining the effects of nimesulide also examined the effects of meloxicam: of 12 healthy dogs *Canis lupus familiaris* treated with standard daily doses over 10 days, those given nimesulide showed evidence of renal damage while those given meloxicam did not (Borges *et al.* 2013). Among all the studies examining the effect of meloxicam, 79% reported a positive effect (e.g. decreased pain level, increased recovery rate, no complications), 9% a mixed effect, 3% a negative effect, and 9% reported no effect (SAVE 2016). The paucity of published evidence for the efficacy and safety of nimesulide as a veterinary drug should raise concern over its use.

It is important to note that this study was funded and carried out through a collaboration of conservation non-government organisations and academic institutions; in the future, the responsibility for testing the impacts of drugs on non-target animals should fall on the pharmaceutical industry, and be carried out *before* the drugs are licensed for veterinary use. Furthermore, we call for a comprehensive ban on the manufacture, distribution, retail and use of all forms of nimesulide in South Asia, except for single-dose vials of human formulations. This will not affect the welfare of domesticated animals as a safer and more effective alternative, meloxicam, is already common and widely available. South Asia’s critically endangered *Gyps* vultures will not recover and the actions to date to rid the region of diclofenac will be severely undermined if treatment of domesticated animals with nimesulide is allowed to continue.

## CRediT authorship contribution statement

**Toby Galligan:** Conceptualization, Methodology, Formal analysis, Investigation, Data Curation, Writing – Original Draft. **Rhys Green:** Conceptualization, Methodology, Writing – Review & Editing. **Kerri Wolter:** Resources. **Mark Taggart:** Methodology, Formal Analysis, Investigation, Resources, Data Curation, Writing – Review & Editing. **Neil Duncan:** Methodology, Formal Analysis, Investigation, Resources. **John Mallord:** Writing – Review & Editing, Project administration. **Dawn Alderson:** Project administration. **Yuan Li:** Formal analysis, Investigation. **Vinny Naidoo:** Conceptualization, Methodology, Formal analysis, Investigation, Resources, Writing – Original Draft

## Acknowledgements

We thank J. Chipangura, R. Mavunganidze and A. van Wyk for technical support in completing the experiments involving cattle; D. Svobodova for assistance in NSAID extraction and analysis; V. Prakash and C. Bowden for obtaining the nimesulide used in this study. This study was funded by the RSPB Centre for Conversation Science, which in turn is funded by the members of the RSPB. The rescue and rehabilitation work of VulPro is funded by the Columbus Zoo, Colchester Zoo, Dallas Zoo, Detroit Zoological Society, DHL, Braak Trust, Hans Hoheisen Charitable Trust, Lomas Wildlife Protection Trust, Natural Encounters Inc. and The Tusk Trust.

## References

Acharya, R., Cuthbert, R., Baral, H.S. & Shah, K.B. (2009) Rapid population declines of Himalayan Griffon Gyps himalayensis in Upper Mustang, Nepal. Bird Conservation International 19: 99–107.

Acharya, R., Cuthbert, R., Baral, H.S. & Chaudhary, A. (2010) Rapid decline of the Bearded Vulture Gypaetus barbatus in Upper Mustang, Nepal. Forktail 26: 117–120.

Arnold, K.E., Brown, A.R., Ankley, G.T. & Sumpter, J.P. (2014) Medicating the environment: assessing the risks of pharmaceuticals to wildlife and ecosystems. Philosophical Transactions of the Royal Society B 369: 20130569.

Bean, T.G., & Rattner, B.A. (2018) Environmental contaminants of healthcare origin: exposure and potential effects in wildlife. pp. 87–122 in Health Care and Environmental Contaminants. A. Boxall & R. Kookana (Eds). Elsevier.

Bennett, A. & Vila, D. (2000) Nimesulide - a non-steroidal anti-inflammatory drug, a preferential cyclooxygenase-2 inhibitor. Expert Opinion on Pharmacotherapy 1(2): 277–286.

BirdLife International (2017) The IUCN Red List of Threatened Species: e.T22695180A154895845. http://doi.org/10.2305/IUCN.UK.2019-3.RLTS.T22695180A154895845.en. Downloaded on 29 January 2021.

Borges, M., Filho, R.M., Laposy, C.B., Guimaraes-Okamoto, P.T.C., Chaves, M.P., Le Sueur Vieira, A. & Melchert, A. (2013) Nonsteroidal anti-inflammatory therapy: changes on renal function of healthy dogs. Acta Cirurgica Brasileira 28: 842–847.

Chaudhary, A., Subedi, T.R., Giri, J.B., Baral, H.S., Bidari, B., Subedi, H., Chaudhary, B., Chaudhary, I., Paudel, K. & Cuthbert, R.J. (2012) Population trends of Critically Endangered Gyps vultures in the lowlands of Nepal. Bird Conservation International 22: 270–278.

Cuthbert, R., Green, R.E., Ranade, S., Saravanan, S., Pain, D.J., Prakash, V., Cunningham, A.A. (2006) Rapid population declines of Egyptian vulture (Neophron percnopterus) and red- headed vulture (Sarcogyps calvus) in India. Animal Conservation 9: 349–354.

Cuthbert, R.J., Taggart, M.A., Saini, M., Sharma, A., Das, A., Kulkarni, M.D., Deori, P., Ranade, S., Shringarpure, R.N., Galligan, T.H. & Green, R.E. (2016) Continuing mortality of vultures in India associated with illegal veterinary use of diclofenac and a potential threat from nimesulide. Oryx 50: 104–112.

European Medicines Agency (2018) Metacam (Meloxicam) : an overview of Metacam and why it is authorized in the EU. European Medicines Agency, London. EMA/CVMP/259397/2006

Fourie, T., Cromarty, D., Duncan, N., Wolter, K., Naidoo, V. (2015) The safety and pharmacokinetics of carprofen, flunixin and phenylbutazone in the Cape Vulture Gyps coprotheres following oral exposure. PloS One 10: e0141419. https://doi.org/10.1371/journal.pone.0141419

Galligan, T.H., Amano, T., Prakash, V.M., Kulkarni, M., Shringarpure, R., Prakash, N., Ranade, S., Green, R.E., Cuthbert, R.J. (2014) Have population declines in Egyptian vulture and red-headed vulture in India slowed since the 2006 ban on veterinary diclofenac? Bird Conservation International 24: 272–28.

Galligan, T.H., Taggart, M., Cuthbert, R.J., Svobodova, D., Chipangura, J., Alderson, D., Prakash, V., Naidoo, V. (2016) Metabolism of aceclofenac in cattle to vulture-killing diclofenac. Conservation Biology 30: 1122–1127.

Galligan, T.H., Bhusal, K.P., Paudel, K., Chapagain, D., Joshi, A.B., Chaudhary, I.P., Chaudhary, A., Baral, H.S., Cuthbert, R.J., Green, R.E. (2019) Partial recovery of Critically Endangered Gyps vulture populations in Nepal. Bird Conservation International 30: 1–16.

Galligan, T.H., Mallord, J.W., Prakash, V.M., Bhusal, K.P., Sarowar Alam, A.B.M., Anthony, F.M., Dave, R., Dube, A., Shastri, K., Kumar, Y., Prakash, N., Ranade, S., Shringarpure, R., Chapagain, D., Chaudhary, I.P., Joshi, A.B., Paudel, K., Kabir, T., Ahmed, S., … Green, R.E. (2020). Trends in the availability of the vulture-toxic drug, diclofenac, and other NSAIDs in South Asia, as revealed by covert pharmacy surveys. Bird Conservation International http://doi.org/10.1017/S0959270920000477

Green, R., Newton, I., Shultz, S., Cunningham, A., Gilbert, M., Pain, D. & Prakash, V. (2004) Diclofenac Poisoning as a Cause of Vulture Population Declines across the Indian Subcontinent. Journal of Applied Ecology 41: 793–800.

Green, R.E., Taggart, M.A., Senacha, K.R., Raghavan, B., Pain, D.J., Jhala, Y., Cuthbert R. (2007) Rate of decline of the oriental white-backed vulture population in India estimated from a survey of diclofenac residues in carcasses of ungulates. PLoS ONE 2(8): e686. doi:10.1371/journal.pone.0000686

Harr, K.E. (2002) Clinical chemistry of companion avian species: a review. Veterinary Clinical Pathology, 31: 140–151.

Herrero-Villar, M., Velarde, R., Camarero, P.R., Taggart, M.A., Bandeira, V., Fonseca, C., Marco, I. & Mateo, R. (2020) NSAIDs detected in Iberian avian scavengers and carrion after diclofenac registration for veterinary use in Spain. Environmental Pollution 266: 115–157.

Hutchinson, T.H., Madden, J.C., Naidoo, V. & Walker, C.H. (2014) Comparative metabolism as a key driver of wildlife species sensitivity to human and veterinary pharmaceuticals. Philosophical Transactions of The Royal Society B Biological Sciences 369: 20130583. http://dx.doi.org/10.1098/rstb.2013.0583

Lierz, M. (2003) Avian renal disease: pathogenesis, diagnosis, and therapy. The veterinary clinics of North America. Exotic animal practice, 6(1), 29–55.

Meteyer, C. U., Rideout, B. A., Gilbert, M., Shivaprasad, H. L. & Oaks, J. L. (2005) Pathology and proposed pathophysiology of diclofenac poisoning in free-living and experimentally exposed Oriental White-backed Vultures (*Gyps bengalensis)* Journal of Wildlife Diseases 41(4): 707–716.

Moore, R.A., Derry, S., Wiffen, P.J. and Straube, S., 2015. Effects of food on pharmacokinetics of immediate release oral formulations of aspirin, dipyrone, paracetamol and NSAIDs – a systematic review. British Journal of Clinical Pharmacology, 80: 381–388.

Mundy, P.J., Butchart, D., Ledger, J., Piper, S. 1992. The Vultures of Africa. Acorn, Johannesburg, South Africa.

Naidoo, V., Wolter, K., Cuthbert, R. & Duncan, N. (2009) Veterinary diclofenac threatens Africa’s endangered vulture species. Regulatory Toxicology and Pharmacology 53: 205–208.

Naidoo, V., Venter, L., Wolter, K., Taggart, M. & Cuthbert, R. (2010a) The toxicokinetics of ketoprofen in Gyps coprotheres: Toxicity due to zero-order metabolism. Archives of Toxicology. 84: 761–766.

Naidoo, V., Wolter, K., Cromarty, D., Diekmann, M., Duncan, N., Meharg, A.A., Taggart, M.A., Venter, L. & Cuthbert R. (2010b) Toxicity of non-steroidal anti-inflammatory drugs to Gyps vultures: a new threat from ketoprofen. Biology Letters 6: 339–341.

Naidoo, V., Taggart, M., Duncan, N., Wolter, K., Chipangura, J., Green, R. & Galligan, T. (2017) The use of toxicokinetics and exposure studies to show that carprofen in cattle tissue could lead to secondary toxicity and death in wild vultures. Chemosphere 190: 80–89.

Oaks, J.L., Gilbert, M., Virani, M.Z., Watson, R.T., Meteyer, C.U., Rideout, B.A., Shivaprasad, H., Ahmed, S., Chaudhry, M.J.I. & Arshad, M. (2004) Diclofenac residues as the cause of vulture population decline in Pakistan. Nature 427: 630–633.

Pain, D., Bowden, C., Cunningham, A., Cuthbert, R., Das, D., Gilbert, M., Jakati, R., Jhala, Y., Khan, A., Naidoo, V., Oaks, J.L., Parry-Jones, J., Prakash, V., Rahmani, A., Ranade, S., Senacha, R., Saravanan, S., Shah, N., Swan, G. & Green, R. (2008) The race to prevent the extinction of South Asian vultures. Bird Conservation International 18: S30–48.

Paudel, K., Galligan, T., Amano, T., Acharya, R., Chaudhary, A., Baral, H.S., Bhusal, K.P., Chaudhary, I.P., Green, R. & Cuthbert, R. (2015) Population trends in Himalayan Griffon in Upper Mustang, Nepal, before and after the ban on diclofenac. Bird Conservation International. 26: 286–292.

Paudel, K., Bhusal, K., Acharya, R., Chaudhary, A., Baral, H., Chaudhaty, I., Green, R., Cuthbert, R. & Galligan, T. (2016) Is the population trend of the Bearded Vulture Gypaetus barbatus in Upper Mustang, Nepal, shaped by diclofenac? Forktail 32: 54–57.

Prakash, V., Green, R., Pain, D., Ranade, S., Saravanan, S., Prakash, N., Venkitachalam, R., Cuthbert, R., Rahmani, A. & Cunningham, A. (2007) Recent changes in populations of resident Gyps vultures in India. Journal of the Bombay Natural History Society 104: 129–135.

Prakash, V., Galligan, T.H., Chakraborty, S.S., Dave, R., Kulkarni, M.D., Prakash, N., Shringarpure, R.N., Ranade, S.P. & Green, R.E. (2017) Recent changes in populations of Critically Endangered *Gyps* vultures in India. Bird Conservation International 29: 55–70.

Saaristo M., Brodin, T., Balshine, S., Bertram, M.G., Brooks, B.W., Ehlman, S.M., McCallum, E.S., Sih, A., Sundin, J., Wong, B.B.M. & Arnold, K.E. (2018) Direct and indirect effects of chemical contaminants on the behaviour, ecology and evolution of wildlife. Proceedings of the Royal Society of London. Series B: Biological Sciences (London) 285: 20181297. http://dx.doi.org/10.1098/rspb.2018.1297

SAVE (2016) Nimesulide: a vulture-toxic drug. RSPB, Sandy. https://save-vultures.org/wp-content/uploads/2019/06/Nimesulide_summary_Nov2016.pdf

SAVE (2020) Report of the 9th Annual Meeting of the Saving Asia’s Vultures from Extinction partnership. RSPB, Sandy. https://save-vultures.org/wp-content/uploads/2020/03/9th-SAVE-Report.pdf

Sharma, A.K., Saini, M., Singh, S.D., Prakash, V., Das, A., Dasan, B.R., Pandey, S., Bohara, D., Galligan, T.H., Green, R.E., Knopp, D. & Cuthbert, R.J. (2014) Diclofenac is toxic to the steppe eagle *Aquila nipalensis*: widening the diversity of raptors threatened by NSAID misuse in South Asia. Bird Conservation International 24: 282–286

Shultz, S., Baral, H.S., Charman, S., Cunningham, A.A., Das, D., Ghalsasi, D.R., Goudar, M.S., Green, R.E., Jones, A., Nighot, P., Pain, D.J., Prakash, V.. (2004) Diclofenac poisoning is widespread in declining vulture populations across the Indian subcontinent. Proceedings of the Royal Society of London B (Supplement) 271: S458–S460. http://doi.org/10.1098/rsbl.2004.0223

Sturkie, P.D. (1986) Avian Physiology. Springer-Verlag, New York.

Swan, G., Naidoo, V., Cuthbert, R., Green, R.E., Pain, D.J., Swarup, D., Prakash, V., Taggart, M., Bekker, L., Das, D., Diekmann, J., Diekmann, M., Killian, E., Meharg, A., Patra, R.C., Saini, M. & Wolter, K.. (2006a) Removing the threat of diclofenac to critically endangered Asian vultures. PLoS Biology 4: e66. http://doi.org10.1371/journal.pbio.0040066

Suleyman, H., Cadiric, E., Albayrak, A. & Halici, Z. (2008) Nimesulide is a selective COX-2 inhibitory, atypical non-steriodal anti-inflammatory drug. Current Medicinal Chemistry 15, 278–283.

Swan, G.E., Cuthbert, R., Quevedo, M., Green, R.E., Pain, D.J., Bartels, P., Cunningham, A.A., Duncan, N., Meharg, A.A., Oaks, J.L., Parry-Jones, J., Shultz, S., Taggart, M., Verdoom, G. & Wolter, K. (2006b). Toxicity of diclofenac to Gyps vultures. Biology Letters 2: 279–282.

Swarup, D., Patra, R.C., Prakash, V., Cuthbert, R., Das, D., Avari, D.J., Pain, D.J., Green, R.E., Sharma, A.K., Saini, M., Das, D. & Taggart, M. (2007) Safety of meloxicam to critically endangered Gyps vultures and other scavenging birds in India. Animal Conservation 10: 192–198.

Taggart, M.A., Senacha, K.R., Green, R.E., Jhala, Y.V., Raghavan, B., Rahmani, A.R., Cuthbert, R., Pain, D.J. & Meharg, A.A. (2007) Diclofenac residues in carcasses of domestic ungulates available to vultures in India. Environmental International 33: 759–765.

Taggart, M.A., Senacha, K.R., Green, R.E., Cuthbert, R., Jhala, Y.V., Meharg, A.A., Mateo, R. & Pain, D.J. (2009) Analysis of nine NSAIDs in ungulate tissues available to Critically Endangered vultures in India. Environmental Science and Technology 43: 4561–4566.

Ulrika, C. & Wong, B.B.M. (2019) Mate choice in a polluted world: consequences for individuals, populations and communities. Proceedings of the Royal Society of London. Series B: Biological Sciences (London) 374: 20180055 http://dx.doi.org/10.1098/rstb.2018.0055

Wolter, K. Neser, W & Hirschauer, M (2015) Cape vulture Gyps coprotheres captive-breeding protocols Version 2. Vulpro, Hartbeespoort.

Zandvliet, M.M. (2005). Electrocardiography in psittacine birds and ferrets. Seminars in Avian and Exotic Pet Medicine14, 34–51

Zorrilla, I., Martinez, R., Taggart, M.A. & Richards, N. (2014) Suspected flunixin poisoning of a wild Eurasian griffon vulture from Spain. Conservation Biology. 29: 587–592.

