## Supplementary Information for "The non-steroidal anti-inflammatory drug nimesulide kills *Gyps* vultures at concentrations found in the muscle of treated cattle"

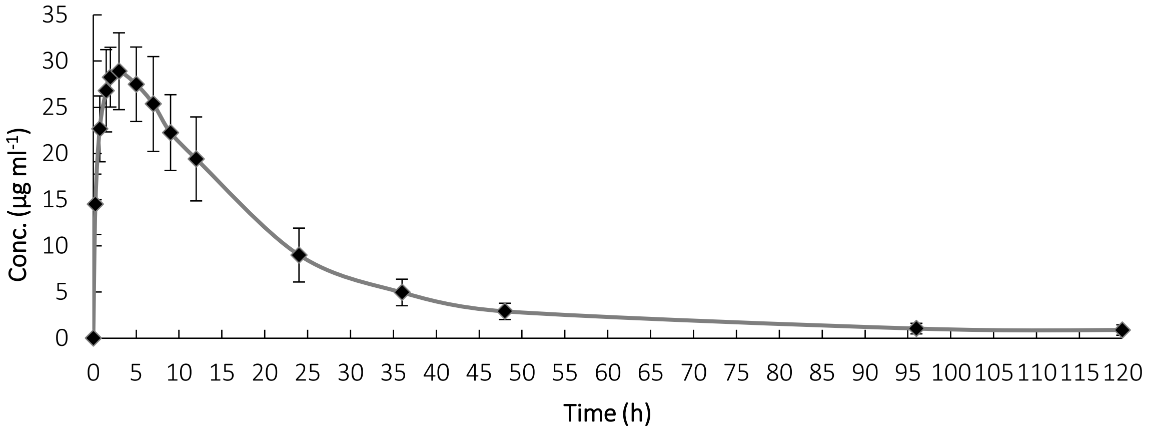


Supplementary Figure 1: The mean plasma concentration (µg/ml) of nimesulide over time (h) in four cattle (C1-C4) treated with nimesulide at 2 mg $kg^{-1}$ bw once. Solid diamonds represent means and vertical lines represent standard deviations.

Supplementary Table 1: Mean concentrations (mg/kg) of nimesulide in three subsamples of three samples of proximal muscle and one sample of three other tissue types harvested from four cattle (C5-C8). The three subsamples were taken from each sample and individually extracted for analysis. Each value for individual cattle represents the mean concentration of the three subsamples. The cattle were treated with nimesulide and slaughtered at the time after dosing when the concentration of nimesulide in plasma was at its greatest ($T_{max}$). The drug was given intramuscularly in the neck. Proximal muscle was harvested from the neck at or close to the injection site and distal muscle was harvested from the hindquarters. Proximal muscle was sampled three times, whereas other tissues were sampled once. The mean concentration of each sample among the four cattle is shown, as is the mean concentration among the mean concentrations of the three samples of proximal muscle.

| Cattle | Proximal muscle 1 | Proximal muscle 2 | Proximal muscle 3 | Distal muscle | Liver | Kidney |
|---|---|---|---|---|---|---|
| C5 | 433.49 | 4.55 | 9.64 | 0.77 | 0.14 | 3.78 |
| C6 | 56.83 | 50.42 | 46.47 | 1.15 | 0.07 | 0.14 |
| C7 | 4.69 | 17.91 | 99.01 | 1.05 | 0.02 | 0.08 |
| C8 | 844.23 | 39.24 | 2.49 | 0.82 | 0.09 | 0.01 |
| Mean (among cattle) | 334.81 | 28.03 | 39.40 | 0.95 | 0.08 | 1.00 |
| Mean (all proximal muscle) | | | 134.08 | | | |

Supplementary Table 2: The individual and mean weights (kg) of four tissue types harvested from four cattle (C5-C8). The cattle were treated with nimesulide and slaughtered at the time after dosing when the concentration of nimesulide in plasma was at its greatest ($T_{max}$). The drug was given intramuscularly in the neck. Proximal muscle was harvested from the neck and distal muscle was harvested from the hindquarters.

| Cattle | Proximal muscle | Distal muscle | Liver | Kidney |
|---|---|---|---|---|
| C5 | 0.740 | 2.168 | 2.756 | 0.552 |
| C6 | 1.529 | 4.255 | 2.696 | 0.628 |
| C7 | 1.150 | 5.164 | 2.630 | 0.428 |
| C8 | 1.030 | 3.706 | 2.493 | 0.561 |
| Mean | 1.112 | 3.820 | 2.640 | 0.540 |
